# The *Caenorhabditis elegans spe-21* gene that encodes a palmitoyltransferase is necessary for spermiogenesis

**DOI:** 10.1101/2025.03.03.641263

**Authors:** Saai Suryanarayanan, Dawn S. Chen, Yamei Zuo, Elizabeth J. Gleason, Zhaorong Zhu, Steven W. L’Hernault, Amber Krauchunas, Andrew Singson

## Abstract

In most animals, spermatids that are produced must further differentiate into fertilization-competent spermatozoa after completing meiosis. In *Caenorhabditis elegans*, this process is known as spermiogenesis or spermatid activation and it results in the transformation of round, non-motile spermatids into amoeboid, motile spermatozoa. Spermatid activation in *C. elegans* is also associated with the fusion of Golgi-derived vesicles called the <u>m</u>embranous <u>o</u>rganelles (MOs) with the plasma membrane. This fusion process is required for producing a fertilization-competent surface on the sperm by placing MO-carried, resident proteins onto the spermatozoon membrane. We have identified, cloned, and characterized the role of *spe-21*, also designated as *dhhc-5,* during both hermaphrodite and male spermatid activation. *spe-21* null mutant worms are severely sub-fertile at all studied growth temperatures. *spe-21* mutant spermatids fail to activate either *in vivo* or *in vitro* after treatment with known chemical activators. We have found that *spe-21* is necessary for MO fusion and pseudopod formation in spermatids. The *spe-21* gene encodes a predicted four pass transmembrane protein with a conserved Asp-His-His-Cys (DHHC) tetrapeptide zinc finger motif embedded in a cysteine rich region (DHHC-CRD type zinc finger motif). Generally, proteins with DHHC-CRD motifs catalyze the post-translational addition of palmitate to their protein substrates and are called palmitoyltransferases or <u>p</u>almitoyl<u>a</u>cyltransferases (PATs). We also found that mNeonGreen-tagged SPE-21 localizes to the MOs in spermatids. Together, our findings show that SPE-21 is an MO localized palmitoyltransferase required for proper spermatid activation and creation of fertilization-competent spermatozoa.

## Introduction

Most animals produce spermatids that must undergo post-meiotic differentiation that is characterized by extensive cytoplasmic and cell surface alteration in the resulting spermatozoa. This process is tightly regulated, and activation happens only when spermatids receive appropriate extracellular cues (Ellis, 2016). In *C. elegans*, spermatid activation, also known as spermiogenesis, occurs either when hermaphrodite-derived spermatids are pushed into the spermatheca during the first few ovulation events or when a male ejaculates into the hermaphrodite uterus (Ellis & Stanfield, 2014). Unlike other animals, gene expression during *C. elegans* spermatogenesis ceases as meiosis is completed (Chu & Shakes, 2013; Ellis & Stanfield, 2014; Han, 2024; L’Hernault, 2009; Miyata et al., 2024; Nishimura & L’Hernault, 2010; Nishimura & L’Hernault, 2017; Ward, 1987) well before spermatid activation. During this process, cellular components required for spermatid activation and fertilization are packaged and sequestered in Golgi derived intracellular, secretory vesicles called the fibrous body-membranous organelles (FB-MOs). MOs first appear in primary spermatocytes as vesicles that bud off from the Golgi. Additionally, cytoskeletal filaments of spermatids that are made of polymeric fibers of major sperm protein (MSP) dimers, form the FBs and begin to associate with MOs (L’Hernault, 2009; Peterson et al., 2021; Price et al., 2021). They are eventually surrounded by double-membrane envelopes derived from the MOs. As secondary spermatocytes divide to form spermatids and a residual body, FB-MOs are sorted into the spermatids (Ward, 1987; Ward et al., 1981; Winter et al., 2017). As spermatids form, the MOs acidify (Gleason et al., 2012), the membrane surrounding the FB retracts into the MO and FB fibers disassemble into constituent MSP dimers (King et al., 1992; Klass & Hirsh, 1981; Nishimura & L’Hernault, 2010; Smith & Ward, 1998) and diffuse throughout the cytoplasm (Price et al., 2021). The MOs localize just beneath the plasma membrane (Q. Wang et al., 2021). During spermatid activation, MOs fuse with the plasma membrane and a pseudopod containing newly, re-polymerized MSP fibers forms (Chu & Shakes, 2013; Peterson et al., 2021) creating motile, amoeboid spermatozoa (Nishimura & L’Hernault, 2010; Nishimura & L’Hernault, 2017; Shimada et al., 2023; Yanagimachi, 1994). MO acidification and fusion are analogous to the acrosome reaction in mammalian sperm (Shimada et al., 2023; Yanagimachi, 1994). Collectively, spermatid activation results in cellular and membrane remodeling that establishes cellular polarity, motility and fertilization competence to spermatozoa.

Several spermatogenesis (*spe*) mutants with defects in meiosis, FB-MO morphogenesis, sperm activation and fertilization have been identified in *C. elegans* (Nishimura & L’Hernault, 2010). One such previously identified mutation that affects FB-MO morphogenesis and sperm activation is in the *spe-10* gene (Gleason et al., 2006; Shakes & Ward, 1989b). In *spe-10* mutant worms, at the end of meiosis, prior to spermatid budding from residual bodies, FB-MOs disassemble, FBs fail to segregate into the spermatids and are retained in the residual bodies. Additionally, the MOs that are retained by spermatids are large, vacuolated and appear like cellular “craters”. These mutant worms also have spermatid activation defects and produce abnormally shortened pseudopods (Gleason et al., 2006; Shakes & Ward, 1989b). The *spe-10* gene encodes a lipidating enzyme, also called a palmitoyltransferase or protein acyltransferase (PAT) that localizes within FB-MOs. SPE-10 is a sperm-specific, transmembrane protein with a zinc finger motif and a conserved Asp-His-His-Cys (DHHC) motif in a cysteine-rich domain (CRD) (Böhm et al., 1997; Gleason et al., 2006). While there are 15 predicted DHHC protein coding genes in *C. elegans*, *spe-10* is the only gene cloned and phenotypically characterized thus far (Edmonds & Morgan, 2014).

In this paper, we describe a second identified *dhhc* gene mutation in *C. elegans*, *spe-21,* also designated as *dhhc-5*. The *spe-21* gene encodes a four-pass transmembrane zinc finger protein with a DHHC-cysteine rich domain motif. Like SPE-10, SPE-21 is also sperm-specific and localizes to the FB-MOs. *spe-21* mutant hermaphrodites and males are both severely sub-fertile at all growth temperatures due to spermatid activation defects. Although *spe-21* and *spe-10* mutants have some similar phenotypes like sterility and mild spermatogenesis defect like a sometimes mis-localized nucleus in spermatids (Gleason et al., 2006), unlike *spe-10* mutants, *spe-21* mutants cannot fuse its MOs with the plasma membrane during spermatid activation and this mutant does not form a pseudopod. In conclusion, our data suggest that SPE-21, presumably through its palmitoyltransferase function, facilitates MO fusion and pseudopod creation during sperm activation which is crucial for producing fertilization competent spermatozoa.

## Materials and methods

### Strain maintenance

Nematode strains used in the current study were generated and maintained as previously described (Brenner, 1974). Strains that were used as controls in this study are Bristol N2, *him-5(e1490)*, BA17: *fem-1(hc17ts)*, JK654: *fem-3(q23).* CB224: *dpy-11(e224)*. The *spe-21* strains used are: *spe-21(as41), spe-21(as41); him-5; asEx98, spe-21(eb99)); him-5; asEx98,* SL1061: *spe-21(hc113),* SL1097: *spe-21(hc113); him-5; asEx98, spe-21(syb4299), spe-21(syb4299); him-5(e1490); asEx98*, *spe-21(syb4179 [spe-21::mNG]), spe-21(syb4179* [*spe-21::mNG*])*; him-5(e1490)* The Hawaiian strain CB4856 for whole genome sequencing mapping. The worms were maintained at 16°C and 20°C and all the experiments were performed at 20°C unless otherwise specified. N2 was used as wild type strain when the strain under analysis was N2 derived. Similarly, if *him-5* was in the background of the strains used, *him-5* animals were used as controls. The *spe-21(hc113)* allele was identified through a forward genetic screen, which was similar to that described in the 1989 paper (L’Hernault et al., 1988) and was the generous gift of Jacob Varkey and Sam Ward. Forward genetic screens in the L’Hernault lab and Singson lab led to the identification of *spe-21(eb99)* (recovered in a noncomplementation screen) and *spe-21(as41)* (recovered in a temperature-sensitive screen) (Singaravelu et al., 2015). These strains were initially maintained by crossing with N2 males and selecting for sterile F2 progeny. With the identification and cloning of the *spe-21* gene through whole-genome sequencing, the PCR rescue array (2 μL of 120 ng/μL) corresponding to the genomic location of *spe-21* was injected together with the injection marker plasmid pPD93_97 (3 μL of 180 ng/μL) (*asEx98*) to maintain all *spe-21* mutant strains.

### F_1_ noncomplementation screen for the identification of spe-21(eb99)

The EMS induced mutation *spe-21(eb99)* was isolated in an F_1_ noncomplementation screen to *spe-21(hc113)*. Wild-type males (N2) were mutagenized with 50 mM ethyl methane sulfonate (EMS) for four hours and then mated with homozygous *spe-21(hc113)* hermaphrodites. Eleven thousand five hundred L4 stage outcross hermaphroditic progeny were picked individually at 25°C and screened for self-sterility associated with oocytes on the growth plate. Candidates were picked and mated to wild-type (N2) males and re-tested by complementation to *spe-21(hc113)*.

### Brood and Ovulation counts

Several L4 hermaphrodites were picked onto individual one-spot plates (n>=10). They were allowed to self-fertilize, and every 24 hours they were transferred onto fresh plates until they stopped laying eggs/oocytes. The total number of progeny produced after the eggs hatched was recorded as brood size. The number of progeny and oocytes that were produced by hermaphrodites throughout its reproductive lifespan was noted as the number of ovulation events. These analyses were performed at 16 °C, 20 °C and 25 °C. The errors are reported as standard error of the means with 95% confidence interval.

### Male fertility analysis

At 25 °C, on several one-spot plates (n>=15), crosses were set up between L4 males and L4 *dpy-11(e224)* hermaphrodite in 4:1 ratio. They were allowed to mate for 24 hours. After 24 hours, the males were removed from the plates, hermaphrodites were allowed to lay eggs overnight, and the progeny were counted the next day. Progeny were scored for non-Dpy progeny (outcross progeny) and Dpy progeny (self-progeny). Unmated *dpy-11(e224)* hermaphrodites were also used as controls. The errors are reported as standard error of the means with 95% confidence interval.

### Light microscopy

Differential interference contrast microscopy (DIC) imaging was performed on live or dissected worms. In case of live worms, they were mounted on 2% agarose pads made in M9 media. The worms were kept immobile by a 1 mM solution of levamisole in M9. DIC images were captured using a Zeiss Universal microscopes equipped with an Optronics camera or a ProgRes camera (Jenoptik). The respective software are Magnafire image software (Karl Storz Industrial – America, Inc. El Segundo, CA) and ProgResCapturePro software.

### DAPI staining

Hermaphrodites of varying ages (day 1-day 4) were used for this experiment. These worms were washed at least three times in M9. They were then fixed in ice cold methanol for 30-60 seconds. After fixation, the worms were further washed in PBS and placed on 2% M9 agarose pads and mounted with VECTASHIELD antifade mounting medium with 1.5 μg/ml DAPI (Vector Laboratories, Burlingame, CA). These worms were then used for Nomarski DIC and fluorescent imaging. The contrast of DAPI images of wild-type and *spe-21* mutant hermaphrodites were adjusted uniformly to clearly visualize the sperm in the hermaphrodite reproductive tract.

### Male sperm motility assay

*fem-1(hc17)* L4 hermaphrodites were raised at 16 °C and larval L1 stage worms were shifted and raised at the restrictive termperature of 25°C. At least 15 restrictively-raised L4 hermaphrodites were mated with young adult control or mutant males in 1:4 ratio at 20°C overnight. After 24 hours, hermaphrodites were mounted and DAPI stained to assess sperm transfer and migration in the hermaphrodite reproductive tract.

### *In vitro* male sperm activation

Males that were in L4 were stage were picked and kept celibate by keeping them isolated from hermaphrodites overnight. The following day, their reproductive tracts were dissected using 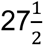 - gauge needles in freshly prepared 1X sperm media (SM) (50 mM HEPES, 25 mM KCl, 45 mM NaCl, 1 mM MgSO_4_, 5 mM CaCl_2_, 10 mM Dextrose; pH 7.8). These dissected sperm served as controls with no activators. To activate spermatids, dissection was done in 1X SM with Pronase E (20 μg/ml) which was allowed to incubate for 7 minutes before the coverslip was placed. Other known activators like triethanolamine (TEA, 120 mM at pH 7.8) and Zinc chloride (20 mM ZnCl_2_ in 1XSM pH 7.0) *in lieu* of Pronase E were also tested for their ability to activate spermatids.

### *In vivo* hermaphrodite sperm activation

L4 hermaphrodites were isolated to avoid any mating with males. As they become young adults, their reproductive tracts were dissected using 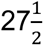 - gauge needles in freshly prepared 1Xsperm media (SM) (50 mM HEPES, 25 mM KCl, 45 mM NaCl, 1 mM MgSO_4_, 5 mM CaCl_2_, 10 mM Dextrose; pH 7.8) with cuts made near spermathecae that releases sperm.

### Whole genome sequencing

Homozygous *spe-21(as41)* hermaphrodites were mated with CB4856 males. The F1 generation of worms were allowed to self-fertilize and about 500 F2 progeny in L4 stage were isolated and picked into 24-well plates. Their fertility was score and approximately 150 F2 self-sterile young hermaphrodites were picked, washed in M9 and collected in Tris-EDTA buffer. These worms were then subjected to genomic DNA extraction using standard column purification method. KAPA Hyper Prep Kit (KAPA Biosystems) was used for library preparation using 10 ng of fragmented DNA. According to Illumina HiSeq instructions, PCR was performed on the DNA library that was generated. Whole genome sequencing of library preparation was performed using an Illumina HiSeq 2500 platform as previously described (Krauchunas et al., 2023) and data analysis was done using MiModD program (https://mimodd.readthedocs.io/en/latest/).

### RT-PCR

Wild-type N2 and *fem-1(hc17)* animals were bleach synchronized and approximately 500-600 adult animals from each strain were collected for RNA extraction using the ZYMO Research Direct-zol™ RNA MicroPrep kit. A cDNA library was constructed from 1 μg of total RNA using the Bio-Rad iScript™ Select cDNA Synthesis Kit (Cat #1708896). PCR was performed using the cDNA as template and *beta-actin* (control) and *spe-21* specific primers were used to assess mRNA expression levels in these strains.

### Northern blot analysis

Total RNA was isolated from *fem-3(q23 gf)* and *fem-1(hc17)* worms. Approximately 20 μg of total RNA was loaded in each lane on a 1.4% formaldehyde agarose gel, subjected to electrophoresis and transferred to a nylon membrane. The ribosomal bands for all the lanes showed that loadings appeared to be equivalent. The resulting blot was hybridized to an 848 bp radiolabeled genomic PCR product, amplified using primers for *spe-21*.

### Transgenic rescue line

The R13F6.5 genomic DNA region specific to *spe-21* was PCR amplified from N2 genomic DNA. Expand Long Template PCR system from ROCHE was used for the same. The PCR product (2 μL of 120 ng/μL) along with marker plasmid pPD93_97 (3 μL of 180 ng/μL) injected into young adult N2 hermaphrodites and selected for stable transgenic lines. When the transgenic line was crossed into the *spe-21* mutant background, GFP-negative F2 worms were homozygous mutants for *spe-21.* The GFP-positive F2 animals from this cross were self-fertile and maintained as transgenic rescue lines.

### *CRISPR/Cas9* genome editing for generating *spe-21(syb4299)*

We collaborated with Suny Biotech (15F/IFC, 1 Wang Long Er Rd, Tai Jiang District, Fuzhou, Fujian, China 350004) to generate all CRISPR/Cas9 genome-edited animals used in the current study. To generate the *spe-21* null allele, s*pe-21(syb4299)*, a stop codon was introduced immediately after the endogenous start codon and immediately following the insertion of a stop codon, deletion of a part of the first exon was also made. This was done to ensure frameshift in the putative null mutant. The genomic status of the null mutant was both PCR and Sanger sequencing-verified. Additionally, these mutants were backcrossed with N2 for three generations to eliminate any potential off-target effects. See supplemental materials and methods for sequence details.

### *CRISPR/Cas9* genome editing for generating *spe-21(syb4179)*

The *spe-21::mNG* strain (*spe-21(syb4179)*) was generated by inserting an in-frame mNeonGreen sequence at the C-terminus of *spe-21* just before the endogenous stop codon. To make the repair site resistant to re-cutting by the Cas9 system, either the PAM site or sgRNA near PAM sites were also mutated to cause synonymous amino acid changes. The animals thus generated were also PCR and Sanger sequencing-verified and backcrossed to N2 for three generations to eliminate any off-target effects. Furthermore, the fertility levels were also verified. See supplemental materials and methods for sequence details.

### MO fusion assay

Spermatids were dissected from *spe-21(syb4299); him-5(e1490)* and *him-5(e1490)* male in 1X SM with Pronase E (20 μg/ml) and FM 1-43 (20 ug/mL). The spermatids stayed in the solution for 7 minutes before putting a coverslip on the slide to ensure complete activation. The activated spermatozoa were imaged using Zeiss Elyra7-Lattice Structured Illumination Microscope (SIM2) with the 63X water objective using the 561 nm laser.

### Localization analysis of SPE-21

#### Spermatids

To visualize the localization of SPE-21 in unactivated spermatids, *spe-21::mNG; him-5* male germline was dissected in freshly made 1X sperm media. The images were obtained on a Leica TCS SP8 tauSTED 3x point scanning confocal microscope under a 40x oil immersion objective and 4X optical zoom. The 510 nm excitation laser was set at 15% intensity, and the emission range was 519-595 nm. This was done with time gating set between 0.3 ns and 6 ns. Standard lightning deconvolution was applied.

#### Spermatozoa

To visualize the localization of SPE-21 in spermatozoa, we acquired live-cell confocal images using a Leica SP8 TCS SP8 tauSTED 3x point scanning confocal microscope system with a 40X mag, 1.30NA oil-immersion objective. First, *spe-21::mNG; him-5* male derived spermatids were activated in vivo. For this experiment, at least 10 *spe-21::mNG; him-5* hermaphrodites were picked and mated with at least 20 *spe-21::mNG; him-5* males overnight. The next day, hermaphrodite reproductive tracts were dissected in freshly prepared 1X sperm media. The 510 nm excitation laser was set at 25% intensity for image acquisition. The emission range was 519 nm-595 nm. This was done along with time gating set between 0.3 ns and 6 ns. The images were acquired with 4X optical zoom. Standard lightning deconvolution was applied.

### SPE-21 sub-cellular localization analysis in spermatids

To identify the subcellular localization of SPE-21 in spermatids, *spe-21::mNG; him-5* male derived spermatids were immunostained with the monoclonal mouse antibody for 1CB4 that has been shown to specifically label the MOs (Okamoto & Thomson, 1985). Spermatids from at least 20 males were dissected in 1X sperm media (SM). These cells were allowed to settle for 5 minutes and then were fixed for 30 minutes in 4% paraformaldehyde in SM, permeabilized for 5 minutes in 0.5% Triton-X in PBS and blocked for 15 minutes in 20% Normal Goat Serum in PBS. All these steps were conducted in a humid chamber and between each step 1X PBS wash was done three times for 5 minutes. The cells were then incubated in a humid chamber, overnight at 4°C with 100 μl primary antibody diluted in PBS-mouse monoclonal antibody 1CB4 used in 1:500 dilution. 1:1000 and 1:2000 dilutions also work. This was again washed with 1X PBS three times for 5 minutes. The samples were then incubated in a humid chamber for 2 hours at room temperature with 100 μl secondary antibody diluted in PBS-1:500 cy-3 conjugated goat anti-mouse antibody was used. 1:1000 dilution also works. The slides were then washed three times for 5 minutes with PBS and the cover slip was placed on the slide. The images were then obtained by sequential scanning on a Leica TCS SP8 tauSTED 3x confocal microscope under 40x oil immersion objective. For the green channel, the excitation laser was set at 500 nm at 25% intensity, and the emission range was 508-541nm with time gating set between 0.3 ns and 6 ns. For the red channel, the excitation laser was set at 554 nm, 0.20% intensity and the emission range was 574 nm-698 nm with no time gating. The images were acquired with 4X optical zoom. Standard lightning deconvolution was applied.

### Generating images showing overlap of PAT proteins from different species

X-ray crystallography structures of human zDHHC20 (PDB 6BMN) and zebrafish zfDHHC15 (PDB 6BMS) were retrieved from RCSB protein data bank. The AlphaFold predicted structure of *C. elegans* SPE-21 (AF-Q21981-F1-model-v4) was obtained from the AlphaFold protein structure database. Then using pyMOLv3.0 (The pyMOL Molecular Graphics System, Version 3.0 Schrodinger, LLC), we generated an image showing strong overlap of all the three structures including the predicted positions of coordinated Zinc ions. Clustal Omega was used for showing the active site conservation of SPE-21 in *C. elegans, H. sapiens and M. musculus*.

### Image analysis and statistics

All the statistical analysis (t-test and ANOVA) were performed with Graph Pad Prism. Statistical significance was defined with a p-value of less than 0.0001. For image analysis, ImageJ Fiji was also used. Image J, FIJI (JACoP) analysis was used for quantifying co-localization analysis and measuring Pearson co-efficient. Additionally, Adobe Illustrator and Adobe Photoshop programs were used to build the figures for the paper.

## Results

## *spe-21* mutant hermaphrodites and males are severely sub-fertile due to sperm specific defects

We have three independently isolated recessive *spe-21* mutant alleles*, spe-21(as41ts)* (Singaravelu et al., 2015)*, spe-21(hc113ts)* (Dr. Diane Shakes thesis, 1988) and *spe-21(eb99ts*) (L’Hernault lab) from various mutant screens. Additionally, a putative genetic null allele, *spe-21(syb4299)* was created by CRISPR and it deletes part of exon 1 and introduces several premature stop codons. All four of these alleles fail to complement each other and each self-sterile mutant had restored self-fertility when in trans to an extrachromosomal transgene carrying a wild-type copy of R13F6.5 genomic DNA. R13F6.5 genomic DNA array rescue was used to maintain sterile homozygous mutant worms (Figure 1B).

**Figure 1:**
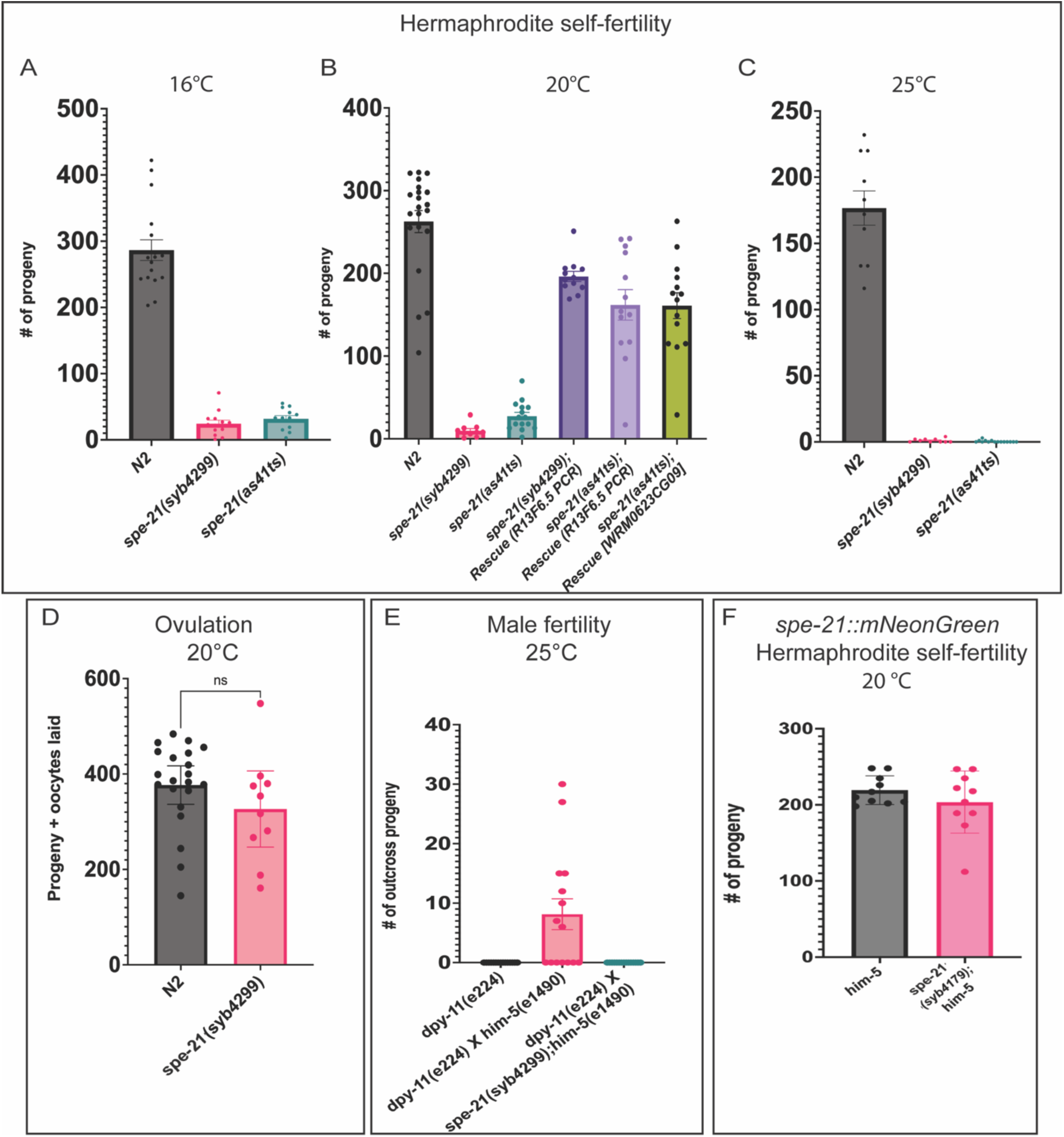
Fertility in *spe-21* mutants. (A-C) Hermaphrodite self-fertility at 16°C, 20°C and 25°C. Self-fertility is measured by the number of self-progeny produced by hermaphrodites. Fosmid WRM0623cG09 or a R13F6.5 templated PCR product recover self-fertility in both *spe-21(as41)* and *spe-21(syb4299)* mutant hermaphrodites. (D) Total ovulation of *spe-21(syb4299)* mutants. This is measure by the total number of progeny, plus the unfertilized oocytes laid per hermaphrodite. For *spe-21(syb4299)* mutants, the majority of ovulations result in unfertilized oocytes. (E) Male fertility at 25°C growth temperature. Males were crossed with *dpy-11* hermaphrodites for 48 hours and number of non-Dpy, outcross progeny were counted in a 24-hour period. (F) Hermaphrodite self-fertility of *spe-21(syb4179)* worms that encode SPE-21::mNG. For all the experiments n ≥ 10 and p-value < 0.0001.

To quantify the effect of *spe-21* mutation on hermaphrodite self-fertility, brood counts of all mutants were performed at 20°C. The *spe-21(as41ts) and spe-21(syb4299)* were also assayed at 16°C and 25°C. *N2* animals were used as wild-type controls. At all temperatures, *spe-21* mutant hermaphrodites were severely sub-fertile (Figures 1A-1C) relative to the N2 wild-type controls. As expected for sperm defective mutants, hermaphrodite cross-fertility was observed after-mating with wild-type males (Supp Figure 1F) (Argon & Ward, 1980; L’Hernault et al., 1988). This suggests that *spe-21* is a gene that shows sperm-specific defects. A Northern blot of fem*-1(hc17)* feminized hermaphrodites and *fem-3(q23)* masculinized hermaphrodites show *spe-21* expression only in the latter population (Supp Figure 2A). Moreover, RTPCR comparing *N2,* and *fem-1(hc17ts)* animals showed bands specific to *spe-21* only in *N2* animals that have sperm (Supp Figure 2B). Together, these results show that *spe-21* expression and function is sperm specific.

**Figure 2:**
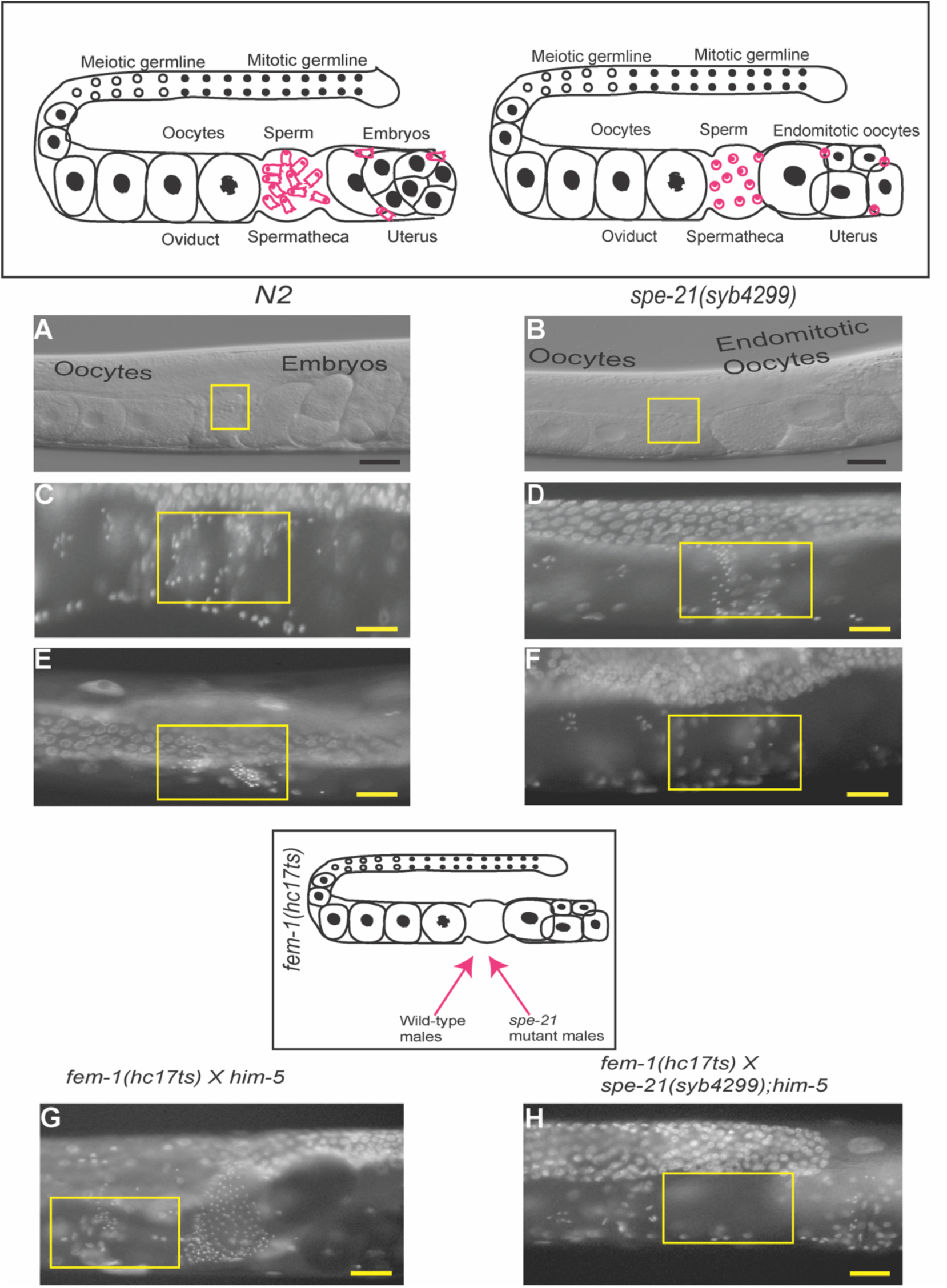
Nomarski DIC and DAPI imaging. DIC images of hermaphrodite reproductive tracts of N2 that have embryos in their uterus (A) and *spe-21(syb4299)* that have only unfertilized oocytes in their uterus (B). Visualizing sperm loss after the start of ovulation in day 1 and day 3 adult *spe-21(syb4299)* (D, F) and *N2* (C, E) worms raised at 20 °C. The positions of the spermathecae are marked by yellow boxes. Sperm nuclei are visualized using DAPI staining as small, bright puncta due the presence of highly condensed nuclear material. (G) *him-5* control males and (H) *spe-21(syb4299)* mutant males mated with *fem-1(hc17)* feminized worms. Wild-type male-derived sperm were able to migrate through the hermaphrodite reproductive tract towards the spermatheca while *spe-21* mutant sperm cannot migrate because they are immotile. Scale bar = 25 µm. Cartoons showing the hermaphrodite reproductive tract are added to this figure to help the readers have a frame of reference.

At 20°C, total ovulation (unfertilized eggs + progeny) levels over the lifespan of *spe-21* mutant animals were measured. The ovulation levels were comparable to wild-type animals (Figure 1D). The minor differences in levels of ovulation between wild-type and mutant animals could be due to faster sperm loss in *spe-21* mutants as it has been previously established that spermatozoa are necessary to stimulate oocyte maturation and ovulation (McCarter et al., 1999; Miller et al., 2001). We observed no significant differences in the brood sizes among the different *spe-21* mutants.

To understand the effect of *spe-21* mutation on male fertility, *spe-21(syb4299); him-5(e1490)* and age matched *him-5(e1490)* males were crossed to *dpy-11* hermaphrodites and the number of non-Dpy, outcross progeny were determined. The number of outcross progeny produced after 48 hours of mating at 25°C in a 24-hour period was significantly different between *him-5(e1490)* and *spe-21(syb4299); him-5(e1490)* males (Figure 1E). Additionally, to test whether male sperm successfully transferred and migrated to hermaphrodite spermatheca, *spe-21(syb4299); him-5(e1490)* and *him-5(e1490)* males were crossed to *fem-1(hc17ts)* hermaphrodites. While *spe-21(syb4299)* sperm successfully transferred to hermaphrodites during mating, they could not migrate to the spermatheca like wild-type sperm and were often found only in ectopic positions like near the vulva and uterus (Figures 2G and 2H). Collectively, these data indicate that *spe-21* mutants produce defective sperm with severely compromised ability to participate in fertilization.

### *spe-21* mutant spermatids cannot maintain their position in the spermatheca and are prematurely depleted

Differential Interference Contrast (DIC) microscopy of adult *spe-21(syb4299)* hermaphrodites raised at 20°C revealed that these animals have normal gonad morphology with oocytes in the oviduct and sperm in the spermatheca. The uterus of *spe-21* mutant hermaphrodites did not have any developing embryos and, instead, were filled with unfertilized, unshelled oocytes (Figures 2A and 2B). DAPI staining was done in age matched N2 and adult *spe-21(syb4299)* hermaphrodites raised at 20°C to visualize sperm position in their germ lines. In DAPI stained worms, sperm due to their highly condensed nature appear as bright puncta. While we did not see significant differences between in the amount of sperm present in day 1 adult *N2* and *spe-21(syb4299)* mutant worms, day 3 adult *spe-21(syb4299)* hermaphrodites had significantly fewer sperm in their spermathecae when compared to wild-type animals (Figures 2C-2F). In wild-type animals, as fertilized oocytes exit the spermatheca and enter the uterus, several sperm are also pushed into the uterus. However, wild-type sperm crawl back into the spermatheca, remain fertilization competent and can go through this cycle repeatedly until they manage to fertilize an oocyte (Kadandale et al., 2005). This suggests that sperm loss in *spe-21* mutants is due to the inability of sperm to efficiently crawl back to the spermatheca. Moreover, when *spe-21(syb4299)* and *him-5* males were crossed to *fem-1(hc17ts)* hermaphrodites, *spe-21(syb4299)* sperm successfully transferred but could not migrate to the spermatheca like wild-type sperm and were often found near the vulva in the uterus (Figures 2G and 2H). This failure to migrate and adhere to spermathecal wall and being displaced by passing oocytes strongly suggest that *spe-21* mutant animals are either defective in spermatid activation and/or spermatozoon motility (Achanzar & Ward, 1997; Singson, 2001; Ward & Carrel, 1979; Zannoni et al., 2003).

### *spe-21* mutant spermatids fail to activate and produce pseudopods *in vivo* and *in vitro* demonstrating the role of *spe-21* in spermiogenesis

To further examine the role of *spe-21* in sperm activation, *in vitro* and *in vivo* spermatid activation assays were performed, and spermatids were examined using Nomarski DIC microscopy. For *in vitro* activation, spermatids were dissected from unmated, young adult *spe-21(syb4299)* and *him-5* males in 1X sperm media alone (Figures 3A and 3C) and further in the presence any of the known *in vitro* activators including Pronase (Ellis & Stanfield, 2014; Nelson & Ward, 1980; Shakes & Ward, 1989a; Ward et al., 1983). While wild-type spermatids produced pseudopods, *spe-21(syb4299)* spermatids failed to activate and did not produce pseudopods or any of the other intermediate cellular extensions like seen in activator-treated *spe-8* class mutants (Krauchunas et al., 2018; Muhlrad et al., 2014) (Figures 3B and 3D) (Shakes & Ward, 1989b). This indicates that *spe-21* mutants have a severe spermatid activation defect. This defect was also observed *in vivo* when sperm were dissected from *spe-21(syb4299)* adult hermaphrodites (Figures 3E and 3F). We also saw that nuclei are eccentrically located in most mutant spermatids (Supp Figure 3A and 3B). This is still a mild spermatogenesis defect as we have observed nuclei off-centeredness in 4% of the wild-type spermatids (Supp Figure 3C) that presumably can still undergo spermiogenesis normally and fertilize the oocytes. Furthermore, *spe-10* mutant spermatids that have a similar off-center nuclei phenotype can still make short pseudopods (Gleason et al., 2006). These results suggest that *spe-21* activity is primarily required for pseudopod formation and spermiogenesis rather than during earlier stages of spermatogenesis.

**Figure 3:**
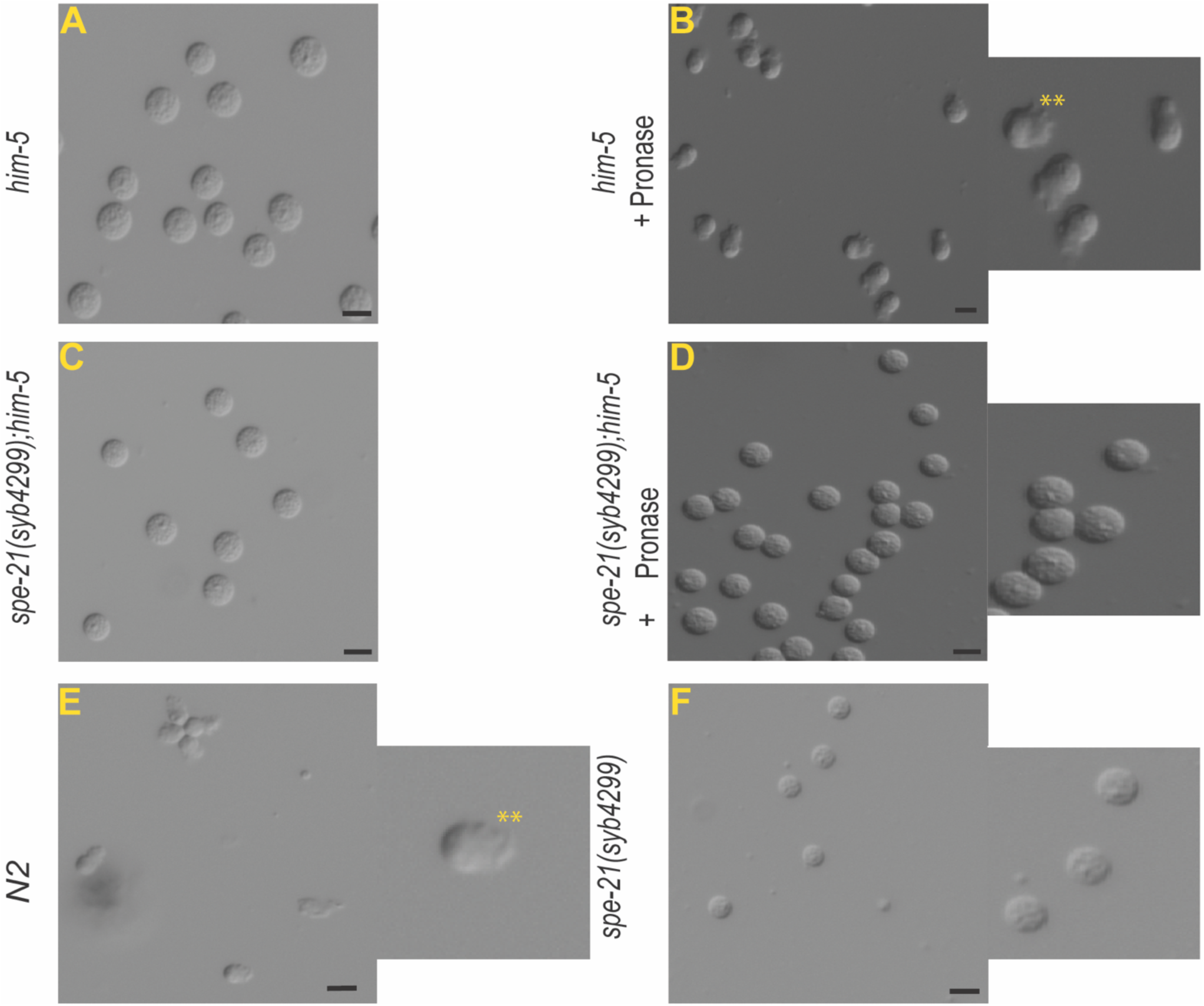
Spermatid activation phenotypes *in vitro* and *in vivo*. (A and B) *him-5* control male-derived spermatids (A) extend pseudopods and become spermatozoa (B) when treated with Pronase E and (C and D) *spe-21(syb4299)* mutant male-derived spermatids (C) fail to produce pseudopods or any intermediate structures (D) after treatment with Pronase E. Dissected *N2* (E) wild-type hermaphrodites release spermatozoa with pseudopods (indicate with **) while dissected *spe-21(syb4299)* (F) mutant hermaphrodites release spermatid-like cells lacking in pseudopods because they are incapable of *in vivo* spermatid activation. Scale bar = 5 µm.

### MOs of *spe-21* mutant sperm fail to fuse with the plasma membrane during sperm activation

MOs are Golgi derived secretory vesicles that begin to form in primary spermatocytes. They are fully mature in spermatids and dock beneath the plasma membrane (PM) of the spermatids (Chu & Shakes, 2013). During spermatid activation, MOs, loaded with cellular components required for sperm activation and fertilization, fuse with the sperm PM. This process results in the delivery of sperm membrane proteins to the PM that are needed to ensure successful fertilization. MO fusion with the PM results in membrane invaginations in the cell body of activated sperm. When these sperm are stained with cell-impermeant lipophilic dyes like FM 1-43, we can observe crescent shaped signal with several bright puncta highlighting the intrusions along the periphery of the cell body (Figure 4) (Washington & Ward, 2006). This occurs because MO fusion allows the dye to enter the now-open MO and stain what had been within its interior. Conversely, in unactivated spermatids, the dye only lights up the outer PM leaflet. We collected spermatids from unmated, young adult *him-5(e1490)* and *spe-21(syb4299); him-5(e1490)* males and treated them with the FM 1-43 dye. In the absence of chemical activators like Pronase E and zinc chloride, in both wild-type and mutant groups, we observed only the PM staining of round spermatids (Figures 4B and 4F). However, in the presence of sperm activators, we observed the following: Wild-type spermatids that activated normally and had MO fusion, showed bright crescent-shaped signals with puncta in the cell body (Figures 4D). This is because the MO fusion pore gives the dye the access to stain the inner membrane. In these spermatids, we also saw pseudopod membrane staining albeit devoid of any bright spots because MO fusion occurs only in the cell body. In *spe-21* mutant spermatids, like the unactivated *him-5(e1490)* control spermatids, we saw only the PM staining implying absence of MO fusion (Figures 4B and 4H). Treating *spe-21(syb4299); him-5(e1490)* spermatids with the activator Pronase E did not change the pattern of FM 1-43 staining or result in the extension of a pseudopod, like it did to the control sperm (Fig. 4D and 4H). These data show that *spe-21* mutants exhibit severe spermiogenesis defects due to a failure of both MO fusion and pseudopodal extension.

**Figure 4:**
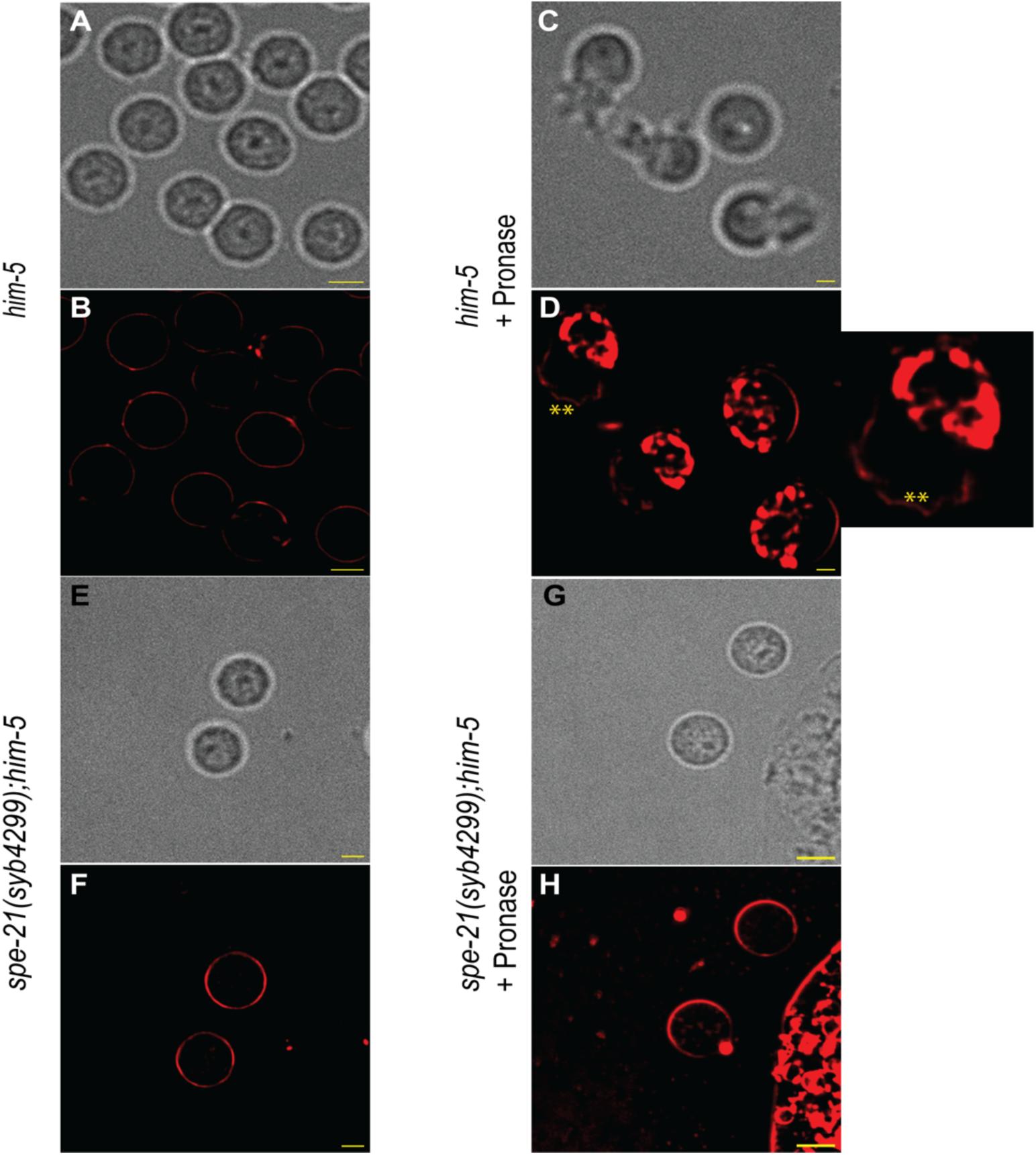
Visualizing MO fusion defects in *spe-21(syb4299)* mutant spermatids. (A-D) Spermatids dissected from *him-5* worms and (E-H) *spe-21(syb4299); him-5(e1490)* mutant worms.(D) Pronase E treated *him-5* control (H) *spe-21(syb4299); him-5(e1490)* mutant male-derived sperm stained with FM1-43 lipophilic dye. MO fusion in the control, activated sperm (D) gives the dye the access to the interior of MOs after MO fusion with the plasma membrane unlike the mutants where only the plasma membrane outer membrane leaflet is stained (H) like the unactivated spermatids dissected from control and mutant animals (B, F). The Asterix symbol (**) in yellow indicates the position of the pseudopods. Scale bar = 5 µm.

### Molecular cloning and identification of *spe-21* gene

Recombination mapping approaches placed *spe-21(hc113ts)* between *lon-1* III and *unc-32* III (Shakes, 1989 Ph.D. thesis). Additionally, with mapping-by-sequencing approach (Doitsidou et al., 2010; Sarin et al., 2008; Thompson et al., 2015; Zhao et al., 2018), we identified a region with low Hawaiian SNPs frequency in *spe-21(as41ts)* mutants. In this region, we found a single missense mutation in Chr:III 6848454 G to A that mapped to the second exon of the gene R13F6.5 (WormBase ID: WBGene00020066). Sanger sequencing also confirmed the presence of this mutation in *spe-21(as41ts)* mutants. Moreover, self-fertility was recovered in *spe-21* mutant hermaphrodites using an extrachromosomal transgene with a wild-type copy of R13F6.5 genomic DNA confirming that *spe-21* indeed is R13F6.5 (Figure 1B).

### *spe-21* encodes a DHHC-CRD zinc finger motif containing palmitoyltransferase

The *spe-21* gene encodes a 244 amino acid containing transmembrane protein (Figure 5B). The predicted SPE-21 protein has four transmembrane domains (TMDs) (Kyte & Doolittle, 1982; Tsirigos et al., 2015), a highly conserved Aspartate-Histidine-Histidine-Cysteine, Cysteine rich (DHHC-CR) zinc finger domain between TMD2 and TMD3 (Böhm et al., 1997; Pagni et al., 2007; Putilina et al., 1999) and a moderately conserved C-terminal palmitoyltransferase C-terminus (PaCCT) motif (Edmonds & Morgan, 2014; González Montoro et al., 2009). Additionally, there are two predicted N-glycosylation sites in the C-terminus (Gupta & Brunak, 2001; Marshall, 1972). The DHHC-CR domain (InterPro entry: IPR001594), C-x_2_-C-x_9_-HC-x_2_-C-x_2_-C-x_4_-DHHC-x_5_-C, is a signature of a distinct family of zinc finger proteins involved in reversible protein S-acylation, commonly known as palmitoylation (Gleason et al., 2006; González Montoro et al., 2009; Lobo et al., 2002; Mitchell et al., 2006; Putilina et al., 1999; Rana et al., 2018; Roth et al., 2002). These proteins are hence referred to as palmitoylacyltransferses (PATs) or palmitoyltransferases (Pfam accession number: PF01529). Furthermore, two Zn^2+^ ions bind the cysteine-rich domain to help maintain the structural integrity of the DHHC active site (Rana et al., 2018).

**Figure 5:**
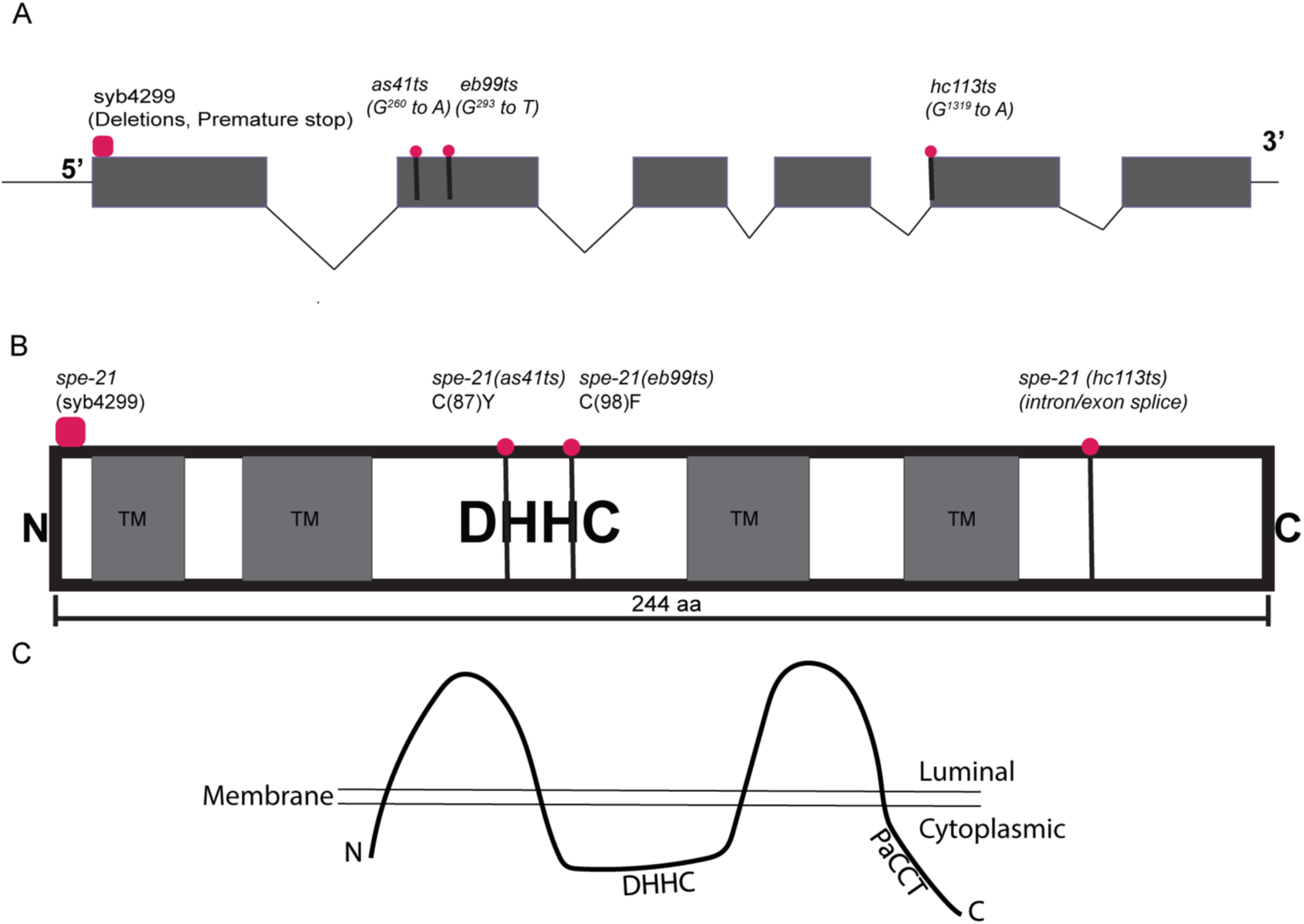
The *spe-21* gene structure, its encoded protein and loss of function mutations. (A) The predicted *spe-21* gene structure with the positions of mutated bases highlighted by magenta circles. (B) *spe-21* encodes a 244 amino acid (AA) protein sequence. The mutated AA residue in each *spe-21* allele is highlighted in magenta. (C) Predicted membrane topology of SPE-21. N and C-termini are both cytoplasmic. The DHHC active site motif is present between the transmembrane domains and is also cytoplasmic.

### Sequences of *spe-21* mutant alleles

There are two known temperature sensitive mutations within the coding region of *spe-21 (*Figure 5A). *spe-21(as41ts)* (Singaravelu et al., 2015) is a G^260^ to A transition resulting in a missense mutation (C87Y) changing the cysteine (C87) in the cysteine rich domain to tyrosine (Y). Similarly, *spe-21(eb99ts)* is a G^293^ to T change causing a missense mutation (C98F) changing the cysteine (C98) in the cysteine rich domain to phenylalanine (F). *spe-21(hc113ts)* is a G^1319^ to A mutation that affects the “acceptor” splice site of the fourth intron (AG to AA) resulting in skipping the fifth exon. Additionally, we generated a null mutant, *spe-21(syb4299*) by deleting regions of the first exon and introducing premature stop codons using CRISPR. Even if mRNA are generated from this locus, the multiple premature stop codons would presumably make this mRNA a target for SMG-mediated degradation (Page et al., 1999; Reeve, 1997). The fertility levels of the temperature sensitive mutants phenocopy the null mutant indicating that all four of the *spe-21* alleles are likely null alleles (Supp Figures 1A-1E).

### SPE-21 localizes to the membranous organelles (MOs) in sperm

To visualize the localization of SPE-21, we generated a tagged allele of *spe-21, spe-21(syb4179)* by inserting an mNeonGreen tag (mNG) to its endogenous C-terminus using CRISPR. The tag does not seem to be interfering with the function of the protein, and this is evident from its hermaphrodite self-brood size being comparable to that of wild-type animals (Figure 1F). We detected signal along the periphery of the cell in spermatids dissected from *spe-21(syb4179)* males (Figure 6D), consistent with MO localization in spermatids. We also immunostained fixed spermatids with the mouse monoclonal antibody, 1CB4, that labels the MOs (Gleason et al., 2006; Okamoto & Thomson, 1985) and observed that the mNeonGreen signal overlapped with the antibody marker (Figures 6G-6J). This co-staining experiment indicates that SPE-21 is a MO resident protein. Additionally, in activated spermatozoa, crescent shaped signal was observed (Figure 6F). This signal was from the plasma membrane of the cell body where MOs fuse during sperm activation and remain attached. Collectively, these results strongly indicate that SPE-21 localizes to the MOs in sperm.

**Figure 6:**
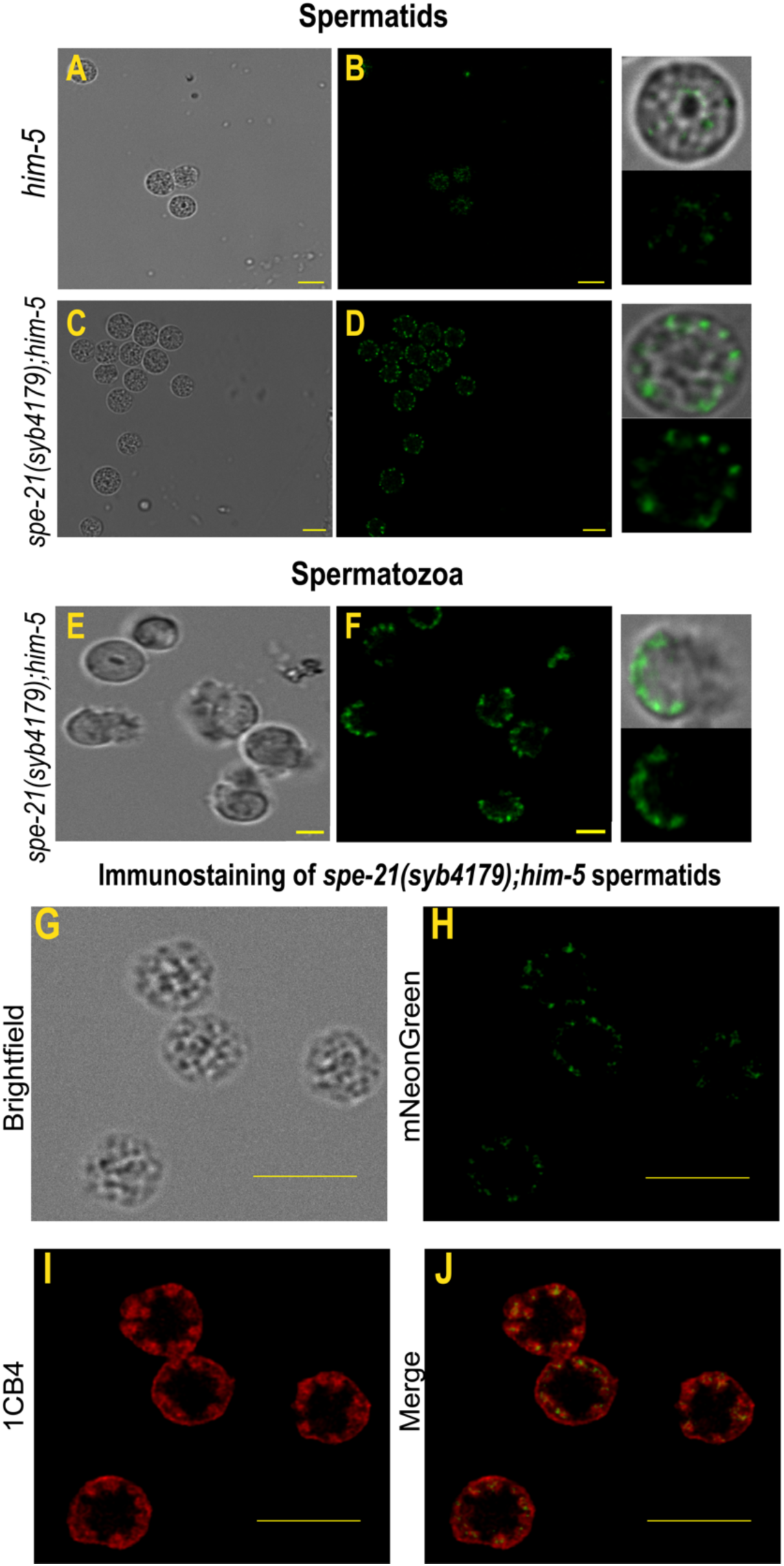
Spermatid and Spermatozoa expressing SPE-21::mNeonGreen protein in live cells. The worm strain labelled as *spe-21(syb4179)* refer to worms that express SPE-21::mNeonGreen. Brightfield images of male-derived unactivated spermatids (A,C) and *in vivo* activated male-derived spermatozoa obtained from *spe21(syb4179)*; *him-5(e1490)* animals (E). We detected a mNeonGreen signal along the periphery of the spermatids dissected from *spe21(syb4179); him-5(e1490)* (D) but not in *him-5(e1490)* (B) spermatids. In *in vivo* activated spermatozoa, a crescent-shaped signal along the cell body of the spermatozoa was seen in *spe21(syb4179); him-5(e1490)* (F). Immunostaining *spe21(syb4179); him-5(e1490)* spermatids with a known MO marker, 1CB4 (G-J) strongly suggests that the SPE-21 is an MO component. Pearson’s coefficient =0.408, Scale bar = 5 µm.

## Discussion

The *spe-21* gene encodes a sperm specific, MO-localized palmitoyltransferase. While there are 15 predicted PAT proteins in *C. elegans*, *spe-21* is only the second characterized gene in this class besides *spe-10* (Edmonds & Morgan, 2014; Gleason et al., 2006). We found that *spe-21* participates in two critical events of spermiogenesis, MO fusion and pseudopod formation which are required to allow a spermatid to become a fertilization-competent spermatozoon. No previously described *spe* mutant that is competent to form spermatids has both of these defects. Among the >60 currently known *spe* mutants, the phenotype that is most similar to *spe-21* is that of *fer-1* mutants in which spermatids fail to fuse their MOs but are able to extend a short pseudopod during spermiogenesis (Washington & Ward, 2006). Future work should reveal how SPE-21 coordinates MO fusion with pseudopod formation in the creation of a functional spermatozoon.

Two of the characterized PAT proteins in *C. elegans*, SPE-10 and SPE-21, are expressed only in sperm, and are components of the MOs (Gleason et al., 2006). Mutations in either *spe-10* or *spe-21* genes cause severe sperm related fertility defects in worms. However, each mutant has a different set of MO-related defects. For instance, MOs in *spe-10* mutants tend to be large and vacuolated but spermatids activate to produce shortened pseudopods. In contrast, *spe-21* does not have any obvious ultrastructural defects in FB-MO structure like vacuolated MOs. Rather, SPE-21 is required for MO fusion with the plasma membrane during spermatid activation. While it is appropriate to hypothesize redundancy between two proteins belonging to the PAT family, the differences in cellular and molecular phenotypes suggest that they have unique and indispensable roles in sperm function. Interestingly, both null mutants, *spe-10(ok1149)* and *spe-21(syb4299)* are sightly leaky with small broods at all growth temperatures (Gleason et al., 2006). Perhaps, this might be due to partial redundancy between SPE-21 and SPE-10. However, further studies with animals that are mutant for both *spe-10* and *spe-21* is required to dissect their independent and collective functions during spermatogenesis.

DHHC-palmitoyltransferases are polytopic, membrane proteins with DHHC motif (Figure 7A) in the cytoplasmic loop (Politis et al., 2005). Previous studies on yeast and human DHHC proteins, showed that these proteins are mostly residents of ER, Golgi or Golgi-derived compartments (Chamberlain & Shipston, 2015; Ohno et al., 2006; Rana et al., 2018). Our current work also establishes that SPE-21 is a component of the Golgi derived MO vesicles. The hydropathy plot of SPE-21 indicates that there are four transmembrane domains with the DHHC-CRD zinc-finger located in the cytoplasm (Figure 5C). The above predicted topology fits well with the mechanism of palmitoyl-CoA attachment to PATs before getting transferred onto the substrate proteins as pools of acyl-CoA are often sequestered in the cytoplasm by acyl-CoA binding proteins (Leventis et al., 1997; Politis et al., 2005). Prior structural studies of DHHC proteins, showed that there are two Zn^2+^ ions binding the cysteine rich region near, but not at the active site cysteine (Myers et al., 1995; Rana et al., 2018). These interactions are considered important to keep the structure of the active site intact (Rana et al., 2018). Both *spe-21(as41ts)* and *spe-21(eb99)* are missense mutations in the cysteine-rich region changing C87 and C98 to tyrosine (Y) and phenylalanine (F) respectively. These mutations phenocopy the null mutant and this could be due to the disruption of cysteine residues that coordinate with Zn^2+^ ions, which would compromise the structure, stability and catalytic activity of the active site (Gottlieb et al., 2015).

**Figure 7:**
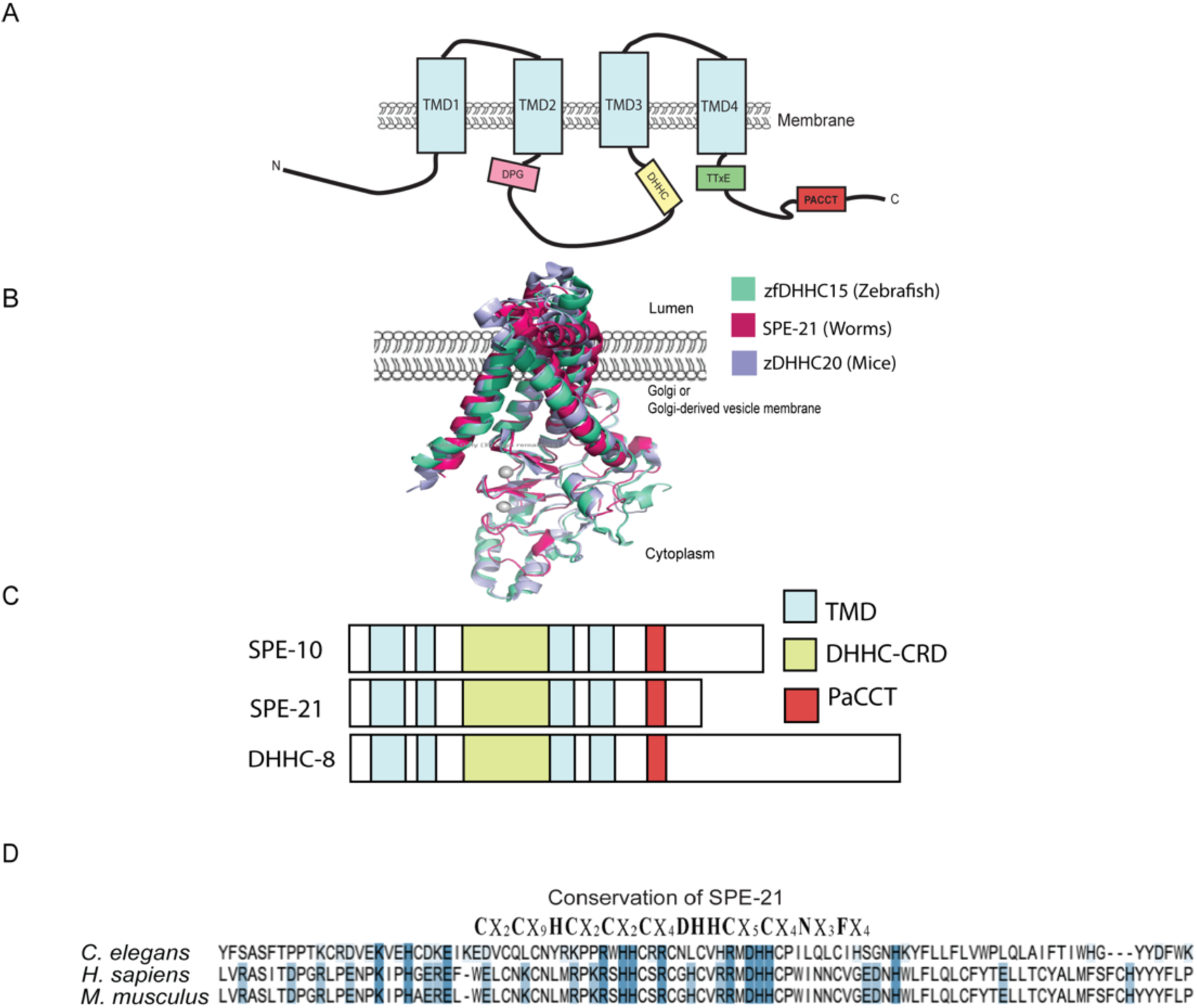
SPE-21 is a highly conserved protein. (A) A schematic showing the membrane topology of SPE-21 and highlighting some of the Pfam motifs present. (B) Using pyMOLv3.0 (The pyMOL Molecular Graphics System, Version 3.0 Schrodinger, LLC) we generated an image showing strong overlap of the X-ray crystal structures of human zDHHC20 (lilac, PDB 6BMN), zebrafish zfDHHC15 (aquamarine, PDB 6BMS) and AlphaFold predicted structure of *C. elegans* SPE-21 (hot pink, AF-Q21981-F1-model-v4). The predicted positions of coordinated Zinc ions are shown as gray spheres. (C) A schematic showing three predicted *C. elegans* DHHC proteins that have a potentially conserved PaCCT motif (red). (D) Clustal omega alignment worm, mice and human SPE-21 protein shows the conservation of SPE-21 active site. All the charged amino acid residues, D, E, K, R and H are highlighted in blue.

DHHC-CRD family of proteins is conserved across different species in both plant and animal kingdoms and there are 15 predicted PATs *in C. elegans* (Edmonds & Morgan, 2014; Fang et al., 2006; Gleason et al., 2006; Hemsley et al., 2005; Mesilaty-Gross et al., 1999; Politis et al., 2005; Rana et al., 2018; Swarthout et al., 2005). Moreover, SPE-21 has homologs in different species including *Homo sapiens and Mus musculus* (Figure 7B and 7D). The DHHC family of proteins are mostly polytopic membrane proteins predicted to have 4 to 6 transmembrane domains (TMDs). Besides the highly conserved DHHC motif, there is substantial diversity in the structure of these proteins. For instance, only some of them possess N terminal ankyrin repeats, a DPG motif (Asp-Pro-Gly) following TMD2, TTxE motif (Thr-Thr-Xaa-Glu) next to TMD4, or a C-terminal PaCCT motif. This variation is perhaps responsible for conferring on PATs their ability to recognize and interact with diverse substrates (González Montoro et al., 2009; Linder & Deschenes, 2003; Lobo et al., 2002; Mitchell et al., 2006; Politis et al., 2005; Rana et al., 2018; Resh, 1999; Roth et al., 2002; Smotrys & Linder, 2004).

Of the 15 predicted PAT proteins in *C. elegans,* only SPE-10, SPE-21 and DHHC-8 are predicted to have a “PaCCT” motif in the C-terminal region (Figure 7C). In the DHHC family of proteins, besides the DHHC-CR domain, the N and C terminal regions are often of variable protein sequence while sometimes containing conserved motifs. This is often attributed to their ability to interact with diverse groups of proteins (González Montoro et al., 2009; Linder & Deschenes, 2003; Lobo et al., 2002; Mitchell et al., 2006; Politis et al., 2005; Rana et al., 2018; Resh, 1999; Roth et al., 2002; Smotrys & Linder, 2004). In yeast, the PaCCT motif was defined as a 16 amino acid region in the C-terminus with an aromatic amino acid in position 3, glycine in position 6, polar residues in position 7, asparagine in position 11 and hydrophobic residues in position 15 and 16 (González Montoro et al., 2009). This region has been hypothesized to be involved in protein-proteins interactions that increases PAT protein stability (Lobo et al., 2002; Swarthout et al., 2005). In SPE-21, we see some of the conserved elements of the PaCCT motif in the 6^th^ exon with tyrosine (Y) position 3, glycine (G) in position 6, serine (S) in position 7, asparagine (N) in position 11 and valine (V) and methionine (M) in positions 15 and 16 respectively. However, among SPE-10, SPE-21, and DHHC-8, there is only moderate conservation further suggesting that their diversity could be important in determining their unique functions and interactions. *spe-21(hc113ts)*, which behaves like the null mutant, is a splice site variant that results in exon skipping by eliminating exon 5. These data suggest that the C-terminal region plays an important role in maintaining the stability, structure, interactions, and function of SPE-21.

Protein palmitoylation has diverse functions across species and cell types. This modification is important for a variety of cellular functions including cell division, cell-cell interactions, protein trafficking and membrane localization, signaling, protein-protein interactions and vesicle fusion (González Montoro et al., 2009; Mesquita et al., 2024). While there is no reported consensus sequence in substrates for palmitoylation, cysteine residues are often the sites of palmitoylation (Reddy et al., 2017). Palmitoylation occurs alone or as an additional reversible lipid modification with prenylation and myristoylation (Resh, 2013). Irrespective of the mechanism, these reversible modifications have physiological relevance in cancers, vascular function, infectious diseases, neural and immune synapse formation (Das et al., 2021; Marin et al., 2016; Pedram et al., 2012; Philippe & Jenkins, 2019; Qian et al., 2022). Interestingly, the fusogenic abilities of vesicle SNARE proteins in neural synapse can be affected by S-acylation. This supports the potential role of SPE-21 in promoting MO fusion with sperm plasma membrane to establish the fertilization synapse. Recent studies have also established the role of zDHHC19, a mouse testis specific PAT protein in spermatogenesis and the acrosome reaction (S. Wang et al., 2021; Wu et al., 2021). Nevertheless, to fully understand the molecular mechanisms involved in germ cell functions and in the establishment of fertilization synapse (Krauchunas et al., 2016), it is important to expand the existing knowledge on lipid modifications in sperm and eggs. Our work shows that the absence of SPE-21 in sperm (Figures 8A and 8B) results in failure of pseudopod formation and MO fusion with the sperm plasma membrane during spermatid activation. Prior work by others has shown that MOs have resident transmembrane and soluble proteins that must translocate to the sperm surface to create a fertilization-competent spermatozoon (Nishimura & L’Hernault, 2017; Singaravelu et al., 2012; Zuo et al., 2023).

**Figure 8:**
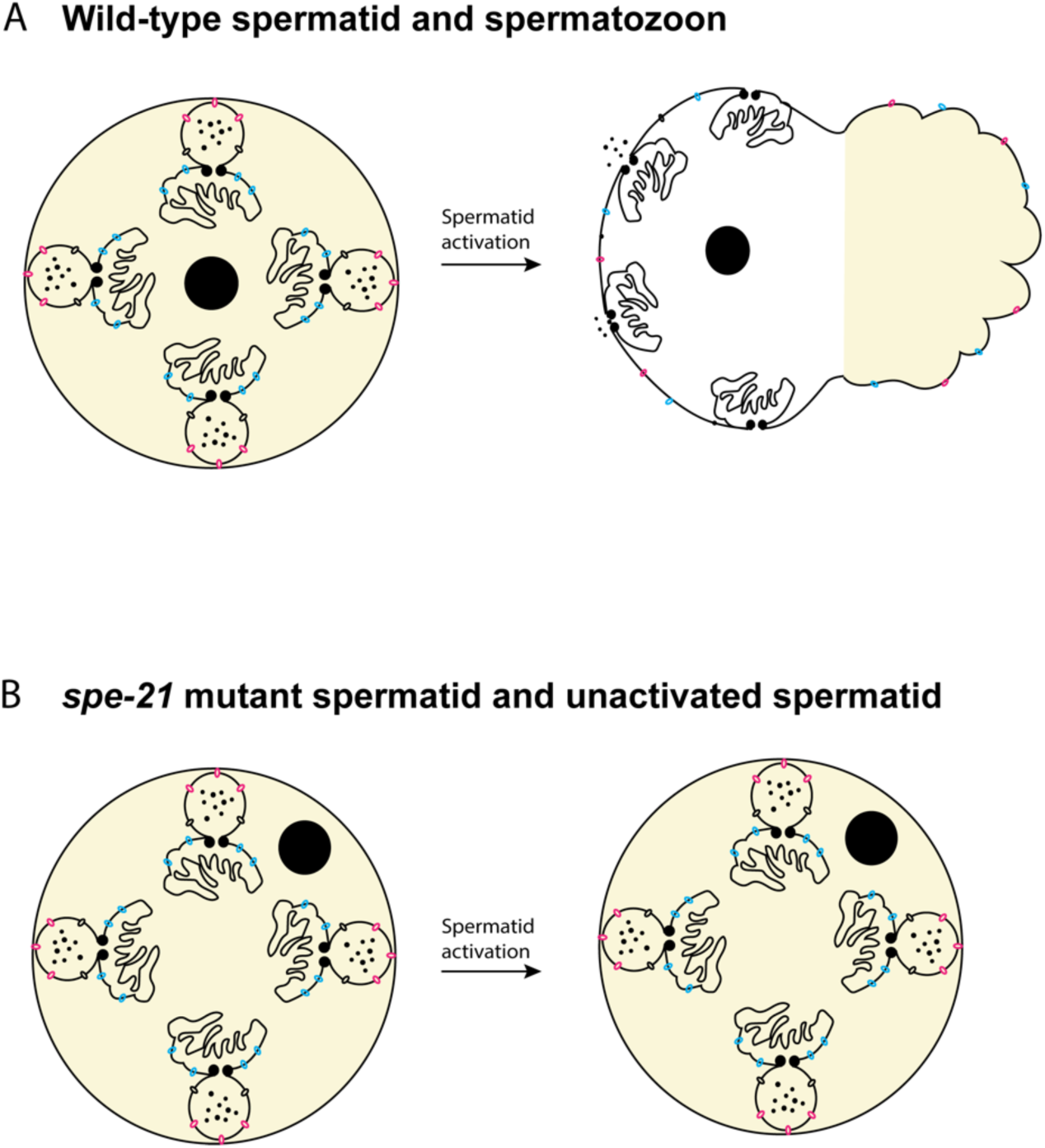
Proposed model for SPE-21 protein function in sperm. In wild-type spermatozoa (A) MOs fuse with the sperm plasma membrane, pseudopods are formed, and transmembrane and soluble proteins get trafficked to the plasma membrane and sperm exterior via MOs. (B) In *spe-21* mutants, MO’s fail to fuse with the plasma membrane and no pseudopod is formed. Beige color is used to represent the distribution of Major Sperm Protein (MSP) distribution in the spermatid and spermatozoon.

In *spe-21* mutants, SPE-21 substrates would likely fail to get palmitoylated and be unable to load onto the MOs and get trafficked to the sperm plasma membrane. As a result, the sperm produced are non-motile and cannot fertilize the oocytes. This shows that SPE-21 plays a critical role in spermatid activation thereby underscoring the significance of lipid modifications in building fertilization-competent sperm.

## Acknowledgements

We thank the imaging core at the Human Genetics Institute of New Jersey and the Waksman Institute Shared Imaging Facility, Rutgers, The State University of New Jersey for the microscopy service. We thank Nanci Kane and Dr. Jessica Shivas for their insights and support with confocal microscopy, instrument training, and data optimization. We would like to thank Dr. Steve L’Hernault, Beth Gleason and Dr. Wes Lindsey, Emory University for generously sharing reagents, worm stains, and any preliminary data with us. We would also like to thank Dr. Diane.C.Shakes, William & Mary and all the members (past and current) of the Singson lab, Waksman Institute, Rutgers, The State University of New Jersey for their invaluable intellectual contributions and insights that greatly enriched this project. We would like to extend our gratitude to Dr. Barth Grant, Dr. Chris Rongo, Dr. Srujana Samhitha Yadavalli and Dr. Karen Schindler for their meaningful discussions and feedback that further enhanced this project. We thank the CGC, funded by NIH Office of Research Infrastructure Programs (P40 OD010440) for worm strains.

## Funding Sources

Research reported in this publication was supported by National Institute of Health (NIH) grants (HD054681) to A.W.S, NIH grant GM40697 to S.W. L. and funds from Emory College, and the Emory University Research Committee award to S.W.L. S.S. was also supported by the Waksman Institute awarded Busch Predoctoral Fellowship and the Acceleration and Completion Fellowships for Rutgers PhD students.

## Author contributions

Conceptualization, S.S., A.S., A.R.K.; Methodology, S.S., A.S., A.R.K., D.S.C., Y.Z., Z.Z., S.W.L., E.J.G.; Investigation, S.S., A.R.K., D.S.C., Y.Z., Z.Z., E.J.G.; Writing – original draft, S.S., A.S.; Writing – review & editing, S.S. A.W.S., A.R.K., S.W.L., E.J.G.; Supervision, S.S., A.S., A.R.K. Funding acquisition, A.W.S., S.W.L. S.S.

## Declaration of interests

No competing interests are declared by the authors.

## Supplementary figures

**Supp Figure 1:**
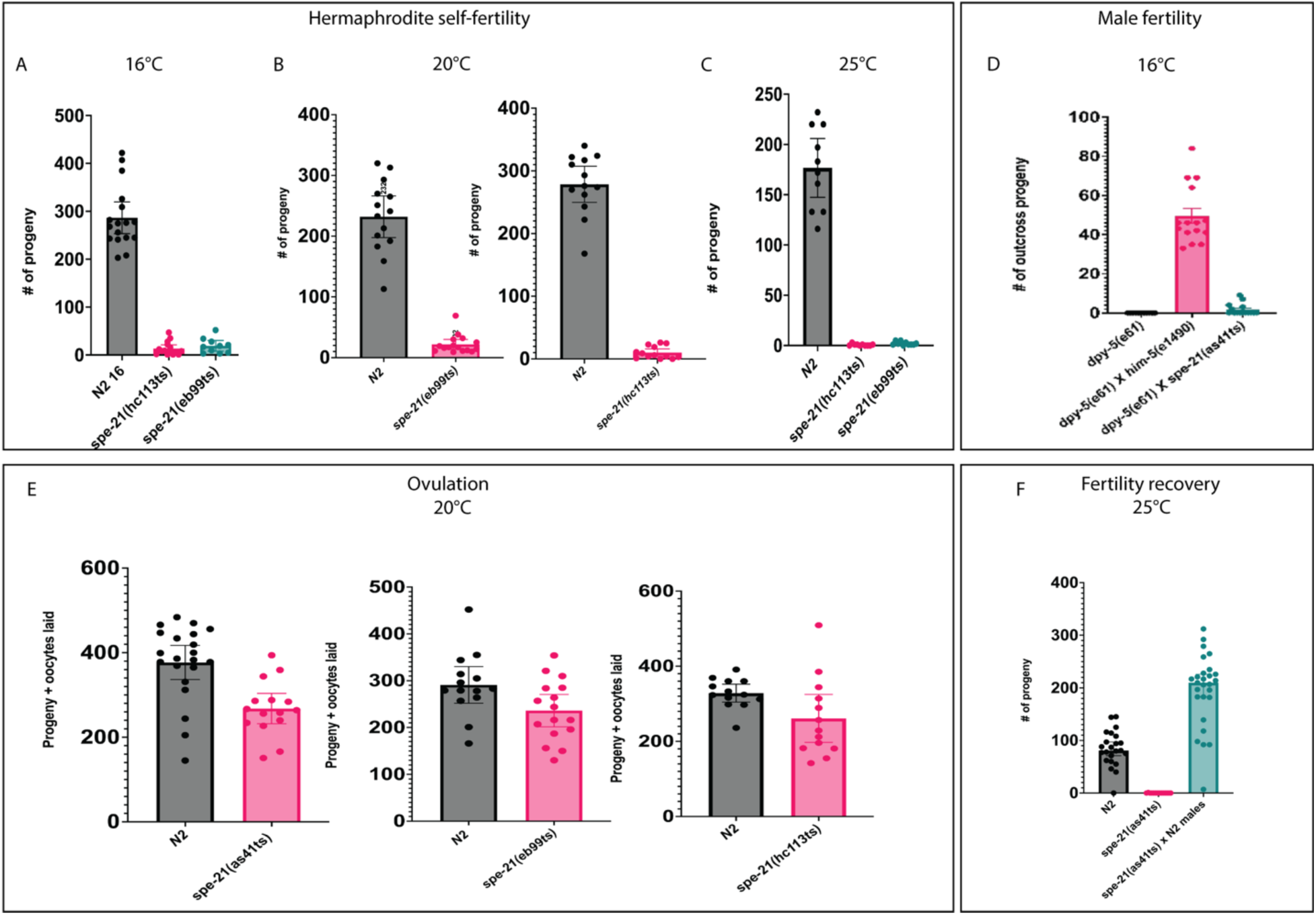
Fertility of *spe-21* mutants. (A-C) Hermaphrodite self-fertility of *spe-21(hc113)* and *spe-21(eb99)* worms grown at 16°C, 20°C and 25°C. The number of self-progeny produced by hermaphrodites is measured as self-fertility. (D) Male cross fertility of *spe-21(as41)* worms grown at 16°C. (E) Total ovulation of *spe-21(as41)*, *spe-21(hc113)* and *spe-21(eb99)* mutants at 20°C. (F) *spe-21(as41)* mutant hermaphrodites exhibited cross-fertility after mating with N2 wild-type males, showing their self-sterility is due to sperm defects. For all the experiments n ≥ 10, p-value < 0.0001.

**Supp Figure 2:**
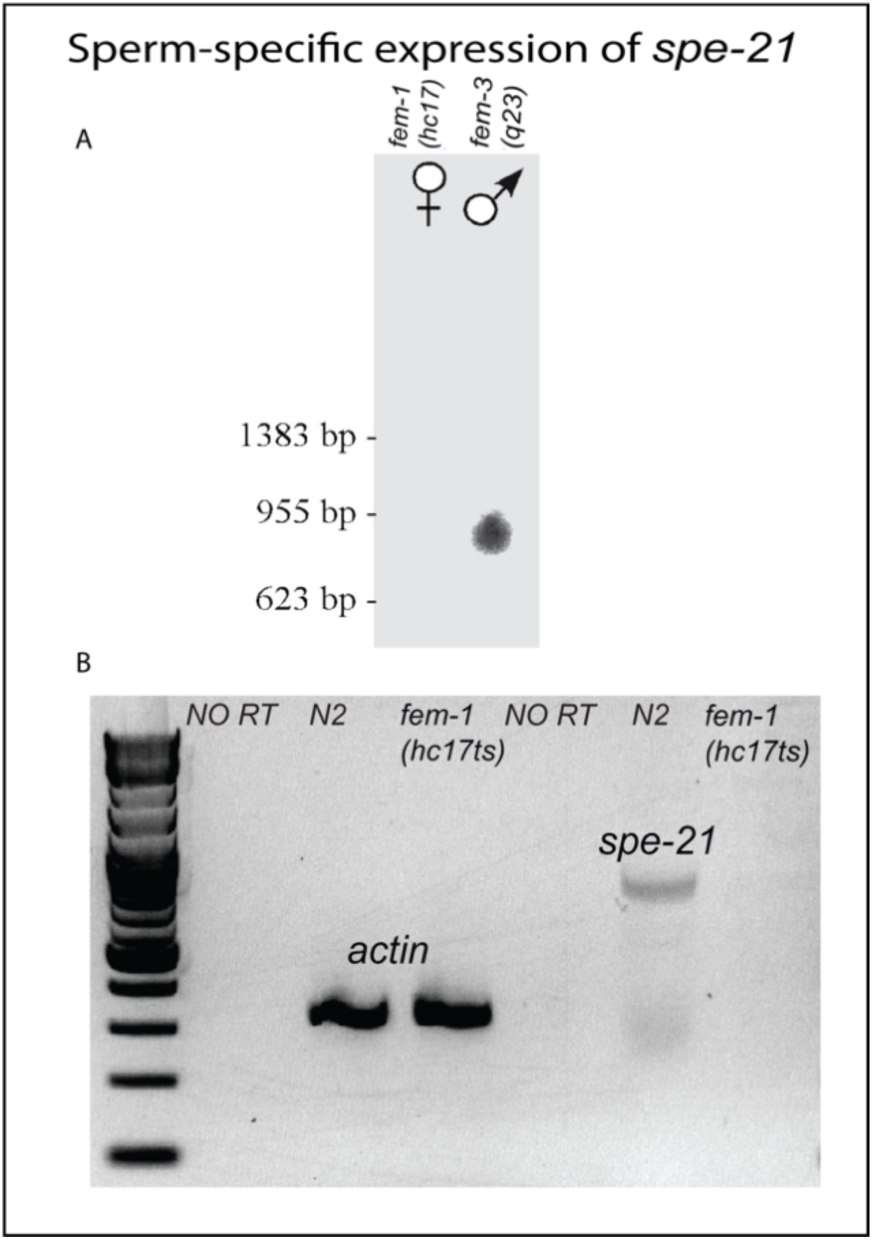
Sperm-specific expression of *spe-21*. (A) Northern blot analysis. RNA extracted from *fem-1(hc17lf),* feminized worms that do not have any sperm (feminized germline) and *fem-3(q23gf)*, worms that do not produce any oocytes and have only sperm (masculinized germline) was run on a gel and the resulting blot was hybridized to an 848 bp long radiolabeled *spe-21* genomic probe. A band for *spe-21* mRNA was only detected in the RNA from masculinized worm. (B) RT-PCR analysis showing a band corresponding to the *spe-21* cDNA only in N2 worms that have both sperm and oocytes. The positive control for this experiment was β actin.

**Supp Figure 3:**
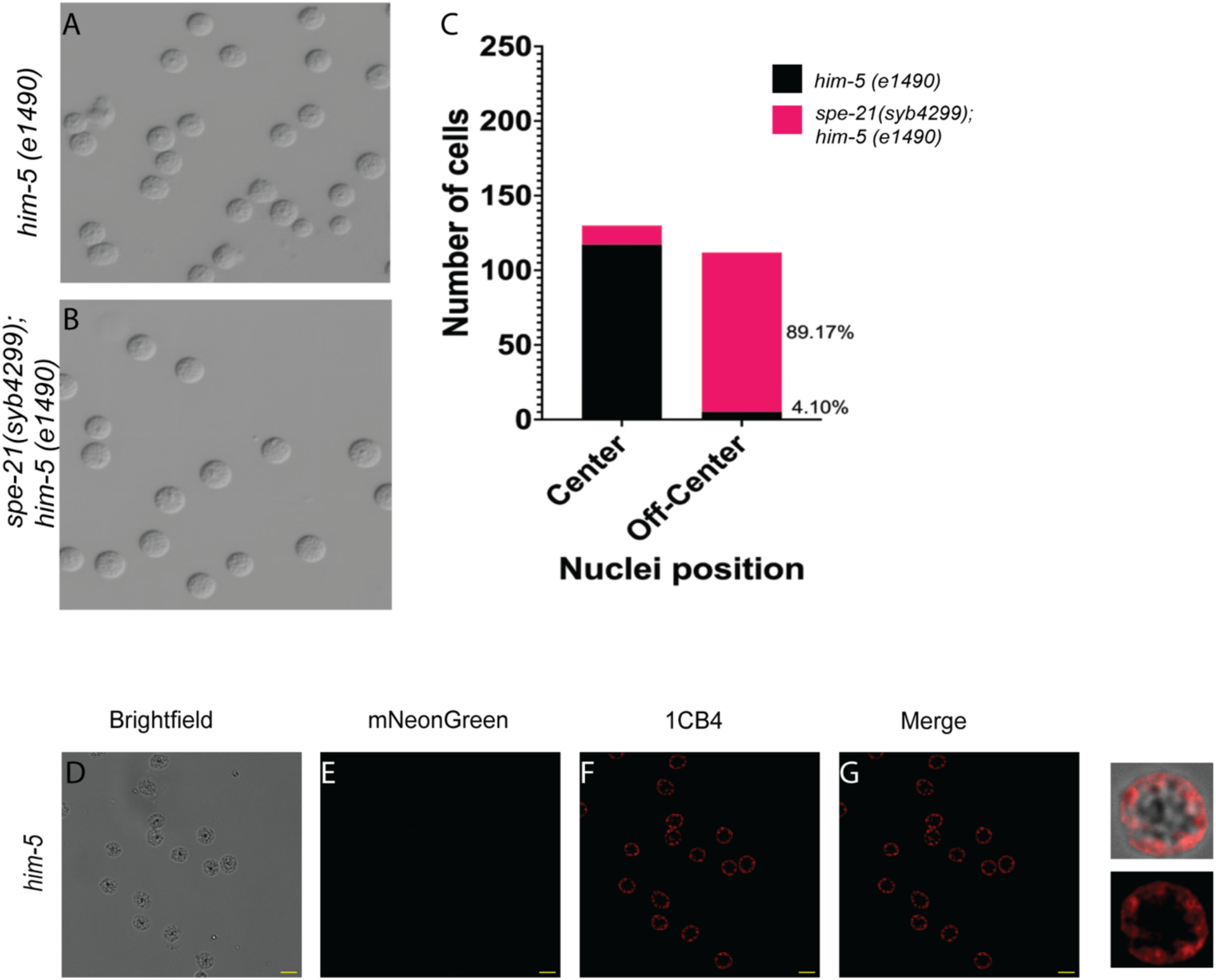
(A, B) Spermatids release from dissected *him-5* and *spe-21(syb4299); him-5(e1490)* mutant males in 1X sperm media. (C) Histogram showing the proportion of cells with off-center nuclei in spermatids from control and *spe-21* mutant worms (D-G) *him-5* spermatids were used as control for the co-localization analysis. The control spermatids were also immunostained like the *spe21(syb4179)* spermatids with a known MO marker, 1CB4. Panel (E) did not have any mNeonGreen signal and panel (F) shows MO staining with 1CB4. Any background signal that might have co-localized with 1CB4 only had the Pearson’s coefficient of 0.088, Scale bar = 5 µm.

## Supplemental Materials and Methods

Sequence information for *CRISPR/Cas9* genome editing for generating *spe-21(syb4299)*:

### Wild-type sequence

cttattatctgattttgccgaATG<u>TGGTTGGAACGCA</u>CACAAAGTGGGTGCTTGCAAACATCGGATGGC

### Sequence deleted

GAACGCA

### Sequence of the final mutant

ATG<u>TGATAT-</u>

CACAAAGTGGGTGCTTGCAAACATCGGATGGCCGATTGCATTCACGTTCTGGTATCAAATCATAGTGGTACTGTACGCAGCTGATGGAACCATCAGTCAATGGGCTCTTTACTACTTTCACTTTCTATGGATTATCATCGTCTGCTCCTATTTTTCGGCATCGTTCACGCCACCGACAAAATGTCGAGATGTTGAGAAGGTGGAACATTGTGATAAGGAAATAAAGGAAGACGTTTGCCAGCTCTGCAACTACCGTAAGCCTCCACGTTGGCATCATTGTCGGAGGTGCAATCTATGTGTTCATCGCATGGATCATCACTGTCCTATTCTGCAGTTGTGCATCCACAGTGGCAATCACAAATATTTTCTACTCTTCCTAGTTTGGCCTCTACAACTTGCAATTTTCACAATTTGGCATGGTTATTACGACTTTTGGAAAACGATAAGAAGCGTTTATACTGCCGAAATTTTAAGCACTTCCGAACAATTGAAGGGAACGGGAGTATCAAATGCACTGATGGTTGGAATTGCAGCTCTGTATCTTCTTAAAAATCAACTTCCAAATCTAATGCGAAATCAGACGTTAATTGAGGAATCAAGGGAGAATACGAGTTACAACCTCGGGTCGTGGCAGGAAAATGTGAAATCAGTAATGGGTGCATGGACTATTGCATGGTTACCTTTTTCAGTTACCAAATCTCGCGAAAAGAAATTCGAATAA

Sequence information for *CRISPR/Cas9* genome editing for generating *spe-21(syb4179)*:

### Wild-type sequence

**<u>ATTGCA</u>**TGGTTACCTTTT**<u>TCA</u>**GTTACCAAATCTCGCGAAAAGAAATTCGAA*TAAttgttatgtcatca ttttattttattgtatagattcaaatcaaaaattctctgaaatttacgtgttcaataaatcat

### Precise mNeonGreen sequence knock-in

**<u>ATCGCT</u>**TGGTTACCTTTT**<u>AGT</u>**GTTACCAAATCTCGCGAAAAGAAATTCGAAATGGTGTCGAA GGGAGAAGAGGATAACATGGCTTCACTCCCAGCTACACACGAACTCCACATCTTCGGATCGATCAACGGAGTGGATTTCGATATGGTCGGACAAGgtaagtttaaacatatatatactaactaaccctgattatttaaattttcagGAACTGGAAACCCAAACGATGGATACGAGGAACTCAACCTCAAGTCGACAAAGGGAGATCTGCAATTCTCGCCATGGATTCTCGTGCCACACATCGGATACGGATTCCACCAATACCTCCCATACCCAGgtaagtttaaactgagttctactaactaacgagtaatatttaaattttcagATGGAATGTCACCATTCCAAGCTGCCATGGTGGATGGATCGGGATACCAAGTTCACCGAACAATGCAATTCGAGGATGGAGCCTCGCTCACAGTGAACTACCGATACACATACGAGGGATCGCACATCAAGgtaagtttaaacagttcggtactaactaaccatacatatttaaattttcagGGAGAGGCTCAAGTTAAGGGAACAGGATTCCCAGCTGATGGACCAGTGATGACAAACTCACTCACAGCTGCTGATTGGTGCCGATCGAAAAAGACATACCCAAATGATAAGACAATCATCTCGACATTCAAGTGGTCGTACACTACTGGAAACGGAAAGCGATACCGATCGACAGCCCGAACAACATACACATTCGCTAAGCCAATGGCCGCCAACTACCTCAAGgtaagtttaaacatgattttactaactaactaatctgatttaaattttcagAATCAACCAATGTACGTGTTCCGAAAGACAGAACTCAAGCACTCAAAGACAGAGCTGAACTTCAAAGAGTGGCAAAAGGCCTTCACAGATGTGATGGGAATGGATGAACTCTACAAGTAAttgttatgtcatcattttattttattgtatagattcaaatcaaaaattctctgaaatttacgtgttcaataaatcat

**Table 1:**
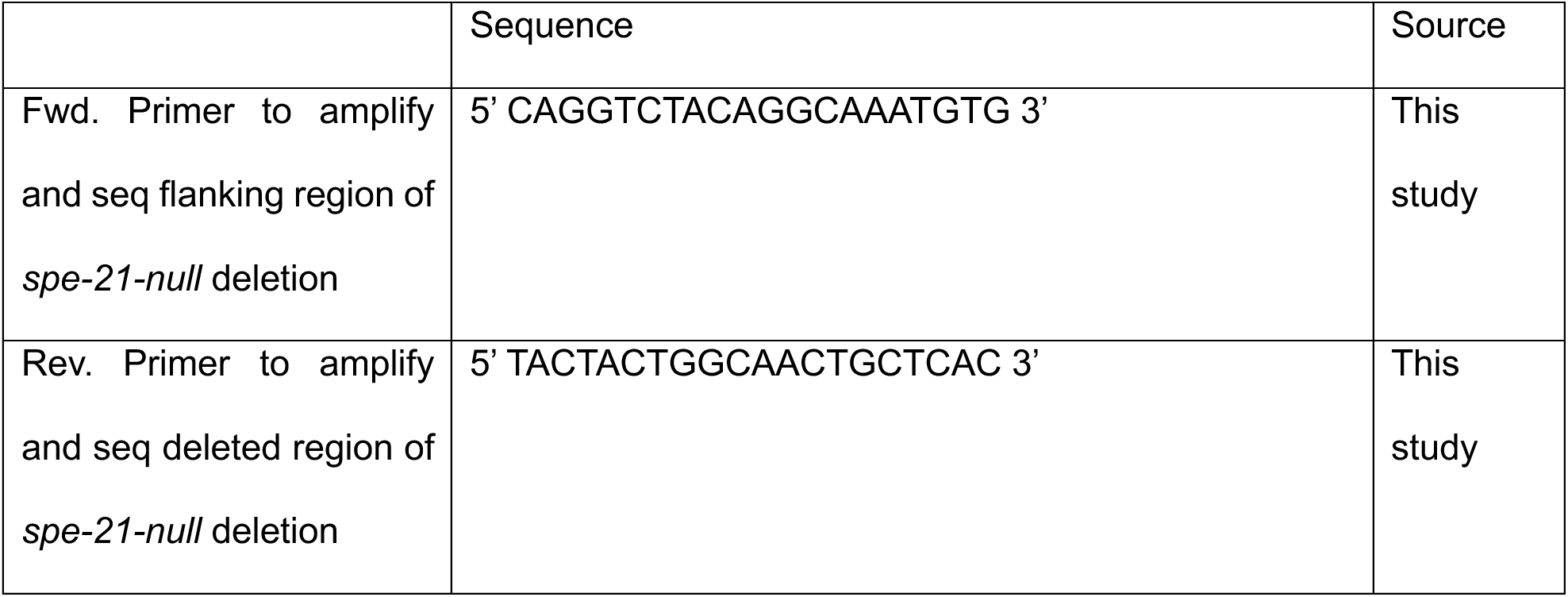
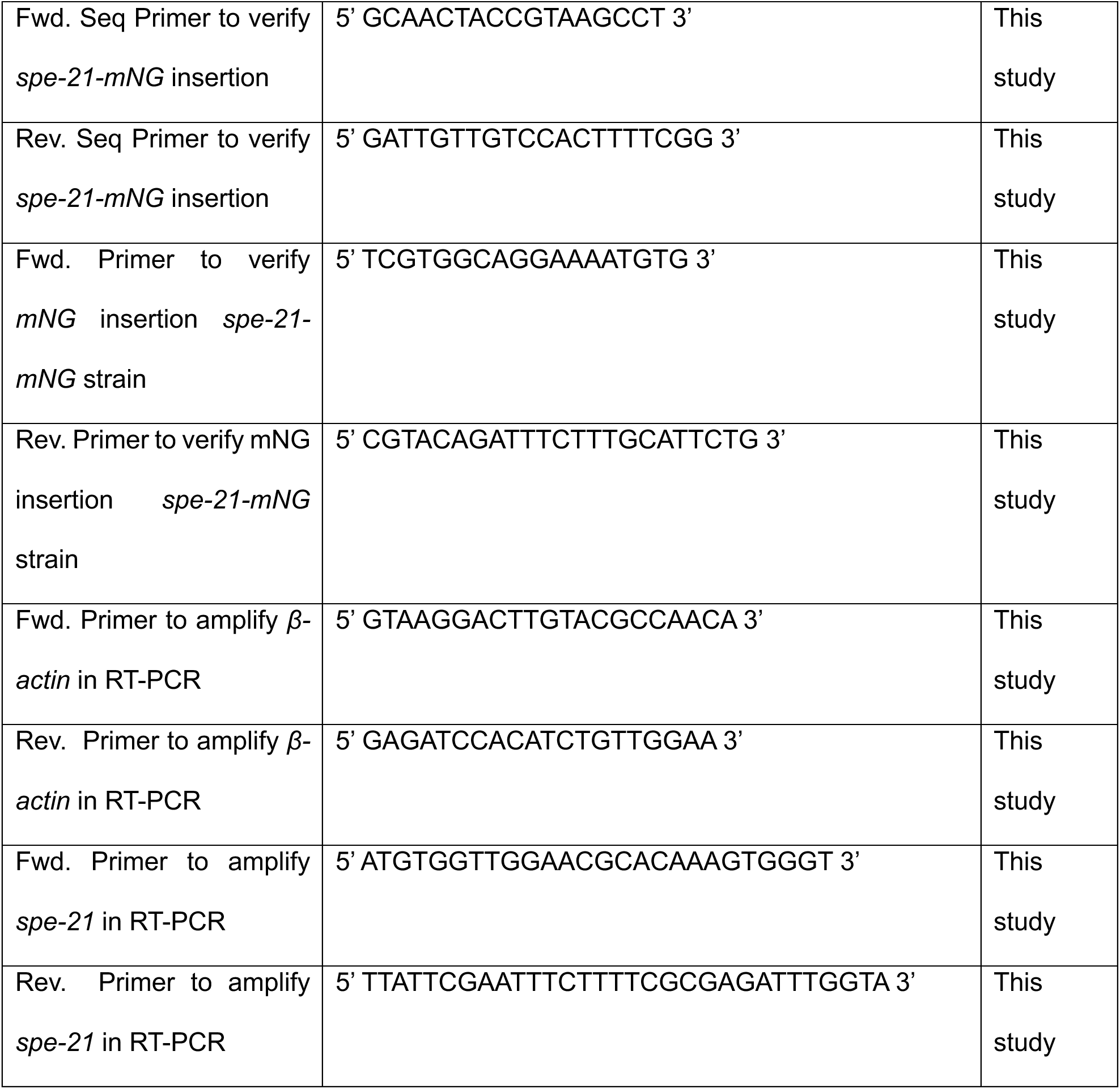
List of primers.

## Notes

### Competing Interest Statement

The authors have declared no competing interest.

